# Deciphering Mechanochemical Influences of Emergent Actomyosin Crosstalk using QCM-D

**DOI:** 10.1101/2024.02.26.582155

**Authors:** Emily M. Kerivan, Victoria N. Amari, William B. Weeks, Leigh H. Hardin, Lyle Tobin, Omayma Y. Al Azzam, Dana N. Reinemann

## Abstract

**Purpose:** Cytoskeletal protein ensembles exhibit emergent mechanics where behavior exhibited in teams is not necessarily the sum of the components’ single molecule properties. In addition, filaments may act as force sensors that distribute feedback and influence motor protein behavior. To understand the design principles of such emergent mechanics, we developed an approach utilizing QCM-D to measure how actomyosin bundles respond mechanically to environmental variables that alter constituent myosin II motor behavior.

**Methods:** QCM-D is used for the first time to probe alterations in actin-myosin bundle viscoelasticity due to changes in skeletal myosin II concentration and motor nucleotide state. Actomyosin bundles were constructed on a gold QCM-D sensor using a microfluidic setup, and frequency and dissipation change measurements were recorded for each component addition to decipher which assay constituents lead to changes in bundle structural compliancy.

**Results:** Lowering myosin concentration is detected as lower shifts in frequency and dissipation, while the relative changes in frequency and dissipation shifts for both the first and second actin additions are relatively similar. Strikingly, buffer washes with different nucleotides (ATP vs. ADP) yielded unique signatures in frequency and dissipation shifts. As myosin II’s ADP-bound state tightly binds actin filaments, we observe an increase in frequency and decrease in dissipation change, indicating a decrease in viscoelasticity, likely due to myosin’s increased affinity for actin, conversion from an active motor to a static crosslinker, and ability to recruit additional actin filaments from the surface, making an overall more rigid sensor coating. However, lowering the ADP concentration results in increased system compliancy, indicating that transient crosslinking and retaining a balance of motor activity perhaps results in a more cooperative and productive force generating system.

**Conclusions:** QCM-D can detect changes in actomyosin viscoelasticity due to molecular-level alterations, such as motor concentration and nucleotide state. These results provide support for actin’s role as a mechanical force-feedback sensor and demonstrate a new approach for deciphering the feedback mechanisms that drive emergent cytoskeletal ensemble crosstalk and intracellular mechanosensing. This approach can be adapted to investigate environmental influences on more complex cytoskeletal ensemble mechanics, including addition of other motors, crosslinkers, and filament types.

---

The cytoskeleton consists of elements such as filaments, crosslinkers, and motor proteins that have vastly different mechanical, motile, and force generating capabilities yet work concertedly to perform large-scale tasks, such as cell division and translocation.^1,11^ In many cases, when the cytoskeleton is studied *in vitro*, investigators are considering one family of the cytoskeleton at a time (e.g. actin and myosin or microtubules and kinesin) and are often working close to the single molecule level.^1,13^ While single molecule studies have revealed invaluable information about motor protein properties such as processivity, stepping, and force generation capability, studies are increasingly finding that new behaviors emerge when motors, proteins, and filaments begin working together in groups that are not the sum of the constituent single molecule properties.^1,3,11^ The design principles behind these emergent cytoskeletal mechanics are not yet understood but will be necessary for understanding the basic rules of life, as well as applications like building synthetic cells, deciphering mechanisms of cytoskeletal-based diseases, and engineering smart materials.^2,3,30,53,60^

Actomyosin emergent mechanics are especially interesting as myosin II behavior is strikingly different between the single molecule level and when in ensembles. As a single molecule, myosin II is not processive, or it does not take multiple steps along an actin filament (AF) before diffusing away, and it has a relatively low force generation capacity of a few piconewtons and a number of different reported step sizes, such as 5 nm, 11 nm, and 30 nm.^1,14,23,32,42,44,48,57^ However, when myosin II works in groups, processivity and force generation increase, though not necessarily in a linear fashion with the number of motors present.^1–3,9,17,22,29,30,38,51,52,56,59,60^ Various approaches have been taken to investigate myosin II ensemble behavior, such as adapting the three-bead optical trapping assay to have multiple motors attached to a bead, adapting the gliding filament assay to measuring unloaded velocity and loaded force generation, or measuring force generation capacity of a thick filament of myosins against a suspended AF.^9,29,30,60^ While these approaches yielded valuable information about behavior deviation between single myosins and those in ensembles, limitations in assay design exist, such as having the myosins attached to a rigid bead or coverslip surface and having their relative orientation fixed due to their attachment. In order to probe how motor self-optimization and AF compliancy affect myosin II ensemble dynamics, Al Azzam *et al.* designed an optical trapping assay to investigate teams of myosin II motors within AF bundles.^2,3^ They found that ensemble myosin II motility and force generation depend on its local environment.^3^ The relative concentration and thus attachment of myosin II motors to the AF bundles change the local stiffness of the AF, dictating force generation capability and step sizes of the myosins.^3^ The effect of myosin II motor activity on AF stiffness has also been demonstrated at the bulk AF mesh level, especially when in concert with other static AF crosslinkers or changes in salt environment.^12,18–20,24,33,39,50,62^ Al Azzam’s results, in conjunction with previous rheological studies of bulk AF systems, suggest that a force-feedback mechanism may exist that is mediated by the AF serving as a mechanical force sensor. Other studies have hypothesized that AFs act as a local and systemic force sensor as well, pointing to emergent changes within the actomyosin cytoskeleton that take place based off of function-specific needs.^12,40,55^ However, the design principles for how these physical signals translate to the molecular actuators and how emergent force-feedback loops are controlled within motor ensembles remain a gap in knowledge.^28,55^

A quartz crystal microbalance with dissipation monitoring (QCM-D) is an instrument that uses a piezoelectric sensor to detect nanogram-level changes in mass, as well as changes in viscoelasticity in systems attached to the sensor surface.^7,10,31^ While QCM-D has traditionally been used to study properties of coatings and polymers, it has started to gain traction in the biophysics field to study changes in mechanical properties of biological materials.^31,49^ A review of QCM-D operation principles can be found by Dixon^10^ and others, and applications for QCM-D in studying the cytoskeleton have been reviewed by Kerivan *et al.*^31^ Most cytoskeletal applications involve studying how cells deform in response to a stimulus or drug. There have been *in vitro* cytoskeletal studies that focus more on binding mechanisms of motors and filaments, as well as gaining a better understanding of how classic *in vitro* motility assays assemble and behave on a non-physiological surface, like a coverslip.^4,5,21,46,54,58^ This is important to understand because methods for constructing model *in vitro* experiments will influence subsequent measurements and thus constrain how well we can correlate *in vitro* experiments to *in vivo* phenomena. However, to our knowledge, no one has used QCM-D to investigate the assembly of hierarchically structured cytoskeletal filament and motor ensembles, as well as monitor changes in system viscoelasticity in real time. In order to better understand how AFs may be acting as a force sensor that drives the force-feedback mechanism between the myosin motors and AFs, we developed an approach using QCM-D to investigate the emergent mechanics of actomyosin ensembles.

A QSense Analyzer QCM-D system from Biolin Scientific and their standard gold sensors were used to gather frequency and dissipation measurements for actomyosin bundle formation (Figure 1). With QCM-D, a decrease in frequency corresponds with the piezoelectric sensor being dampened due to mass addition (Figure 1A).^10,31^ Dissipation monitoring of the QSense system allows for measuring relative viscoelastic changes in samples, denoted by an increase in dissipation when a sample becomes “softer”.^10,31^ To mimic conditions on the gold sensor similarly to what was measured in the optical trapping experiments from Al Azzam *et al*., the actomyosin bundles were built from the bottom up onto the sensor surface. Detailed methods can be found in the supplemental information (SI). First, if not using a new sensor, the gold sensor must be cleaned thoroughly using either a sodium dodecyl sulfate solution and exposure to an ozone chamber, or piranha solution. Sensor cleanliness and age can greatly affect frequency and dissipation measurements if debris is left on the sensor from previous experiments, so care must be taken to maintain high quality sensors. Solutions of poly-l-lysine (PLL 30 *μ*L of 0.1% PLL solution in 30 mL of DI water), actin (200 *μ*L in 1 mL), casein (1 mg in 15 mL), full length skeletal myosin II (0.6 nM), and general actin buffer (GAB, contents) were prepared before measurements and kept on ice until ready to use. As in the previous optical trapping experiments, PLL is used to coat the sensor surface to facilitate AF binding, and casein is used as a blocking agent to prevent non-specific binding of subsequent proteins to the sensor surface.^2,3^ There must be enough volume of each solution made in order to cover the sensor surface and reach a plateau in the frequency and dissipation measurements. With the peristaltic pump set to 0.1 mL/min and anywhere from 10-30 minutes required to reach a plateau in measurement, this required at least 1-3 mL necessary for each component of the assay. This can be somewhat difficult when using expensive protein reagents, such as actin and myosin, to ensure that you have enough protein present for the experiment. After the solutions were prepared, they were added into the QSense system and onto the gold sensor surface using the built-in microfluidic system and peristaltic pump. To achieve a baseline measurement, general actin buffer (GAB) was introduced first, and the frequency and dissipation signals were allowed to stabilize. The initial buffer wash also served as a control in each experiment to demonstrate that adding buffer does not add mass to the surface/change frequency, nor does it change dissipation. Next, the dilute PLL solution was added to coat the sensor surface. We also tried pre-coating the gold sensors with PLL as would be done with coverslips in an optical trapping flow cell assay; however, we found subsequent component addition steps and the frequency and dissipation measurements to be more consistent when adding in the PLL with the pump and letting it settle. Subsequently, the solutions of actin, casein, myosin II, actin, and GAB were added via the pump onto the sensor and allowed to settle over approximately 15-minute periods. We note that we also tried implementing wash steps between the additions of the different reagents (see Figure S1). However, this significantly increased the time of an already lengthy experiment with minimal benefit to obtaining clean frequency and dissipation shift measurements. It is also possible that the pump system could influence the actomyosin distribution on the surface of the sensor. The additions to the sensor surface are in a laminar regime, which is also the case when making flow cells in the optical trapping experiments after which this experiment is modeled; thus, we expect similar distributions of assay components across the sensor. It is also possible to have slight deviations in transitional points within the experiment due to adding the different components and having to switch the pump tubing between solutions for the next infusion.

**Figure 1.**
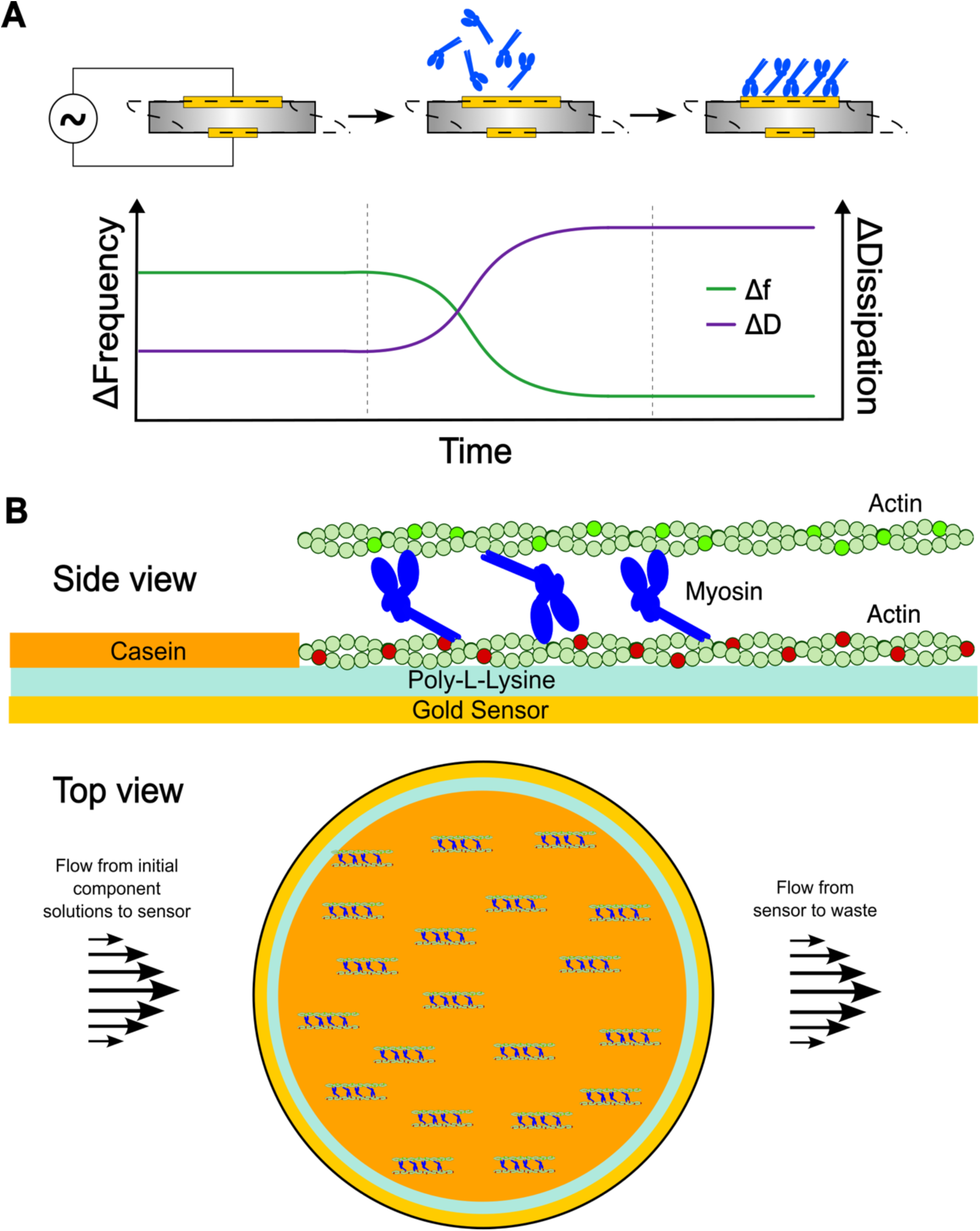
Principles of QCM-D operation and assay schematic. (A) As mass is added to the piezoelectric sensor (vibration indicated through dashed line), frequency decreases (green data, left axis). As viscoelasticity of the sensor “coating” changes from rigid to soft, dissipation increases (purple data, right axis). Adapted from Kerivan *et al.* (2023) *Cytoskeleton*. Reproduced with permission from Wiley under license number 5633140041004. (B) Side and top view of how the actomyosin bundles were assembled onto the gold sensors. After an initial wash with general actin buffer, the following components were introduced via the QSense pump: poly-l-lysine, actin, casein, myosin, actin, and a final buffer wash. Schematics are not drawn to scale.

A representative trace of frequency and dissipation shifts for actomyosin bundle formation is shown in Figure 2. Additional example traces can be found in Figure S1. Several overtone measurements are recorded by the QSense Analyzer with differing balances of signal and sensitivity; overtone 7 exhibits a consistent balance and is used for subsequent analysis. The time to plateau for each component addition varies depending on the concentration of the reagents but is usually around 15 minutes. Again, frequency decrease represents mass addition, and dissipation increase is a change in system viscoelasticity from rigid to more compliant. Changes in frequency and dissipation are minimal for GAB and PLL, as would be expected. The first actin addition settles out relatively quickly in frequency, and a large positive shift in dissipation is apparent, which makes sense as a compliant protein filament is introduced onto the piezoelectric sensor. Casein, the surface blocker, is clearly adding mass, yet the dissipation increase is not as high as for actin since casein is able to evenly and tightly coat the surface. Subsequent additions of myosin and actin show significant changes in frequency and dissipation, especially once actin is added, binds myosin, and makes the final bundle connection onto the sensor surface. A final buffer wash is introduced to ensure that the actomyosin bundles have indeed formed, do not wash away, and serves as a control to demonstrate that introducing more fluid with no proteins to bind the existing bundle does not change the conditions of bundle bound to the sensor surface. These results demonstrate that forming structured, hierarchical cytoskeletal assemblies on a QCM-D sensor is not only feasible, but clear changes in dissipation and thus system viscoelasticity can be observed.

**Figure 2.**
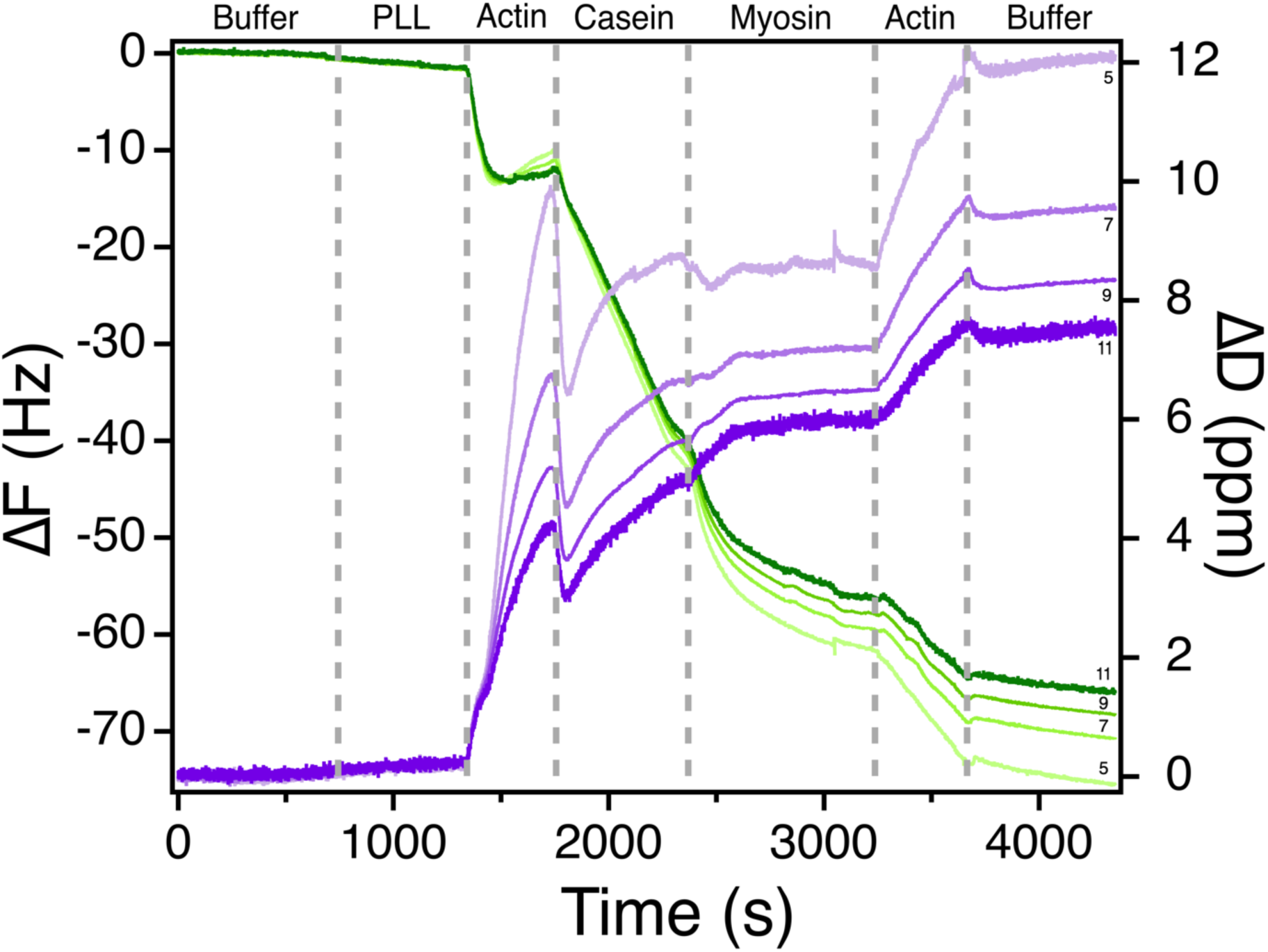
Actomyosin bundle assembly dynamics using QCM-D. Representative QCM-D trace for actomyosin bundle assembly. Change in frequency (green, left axis) and dissipation (purple, right axis) at multiple overtones for each component of the *in vitro* actomyosin bundle assay. These results verify that QCM-D experiments are able to sense molecular-level binding, dynamics, and changes in viscoelasticity of cytoskeletal ensemble systems in real time. Frequency and dissipation shift measurements for specific assay conditions are detailed in Figure 3 and Table 1.

We next wanted to evaluate what types of environmental parameter changes could be measured within actomyosin bundles using QCM-D. As stated earlier, myosin II concentration and activity have been shown to alter systemic force generation in an emergent manner.^3^ As more heads are recruited and bound to AFs, the local environment is less compliant, leading to a sensory-force feedback mechanism of motor activity within the AF bundle.^3^ Thus, we altered myosin concentration to verify that such changes would be measurable with QCM-D. We were also interested in knowing whether changes in small molecules, such as nucleotides, would also be detectable with QCM-D as myosin exhibits a different conformational state and tighter AF binding when ADP is bound than with ATP, thus further altering local system stiffness. Figure 3 demonstrates that lowering myosin concentration from panels A-B to C-D can indeed be detected as lower shifts in frequency and dissipation while the relative changes in frequency and dissipation shifts for both the first and second actin additions are relatively similar.

**Figure 3.**
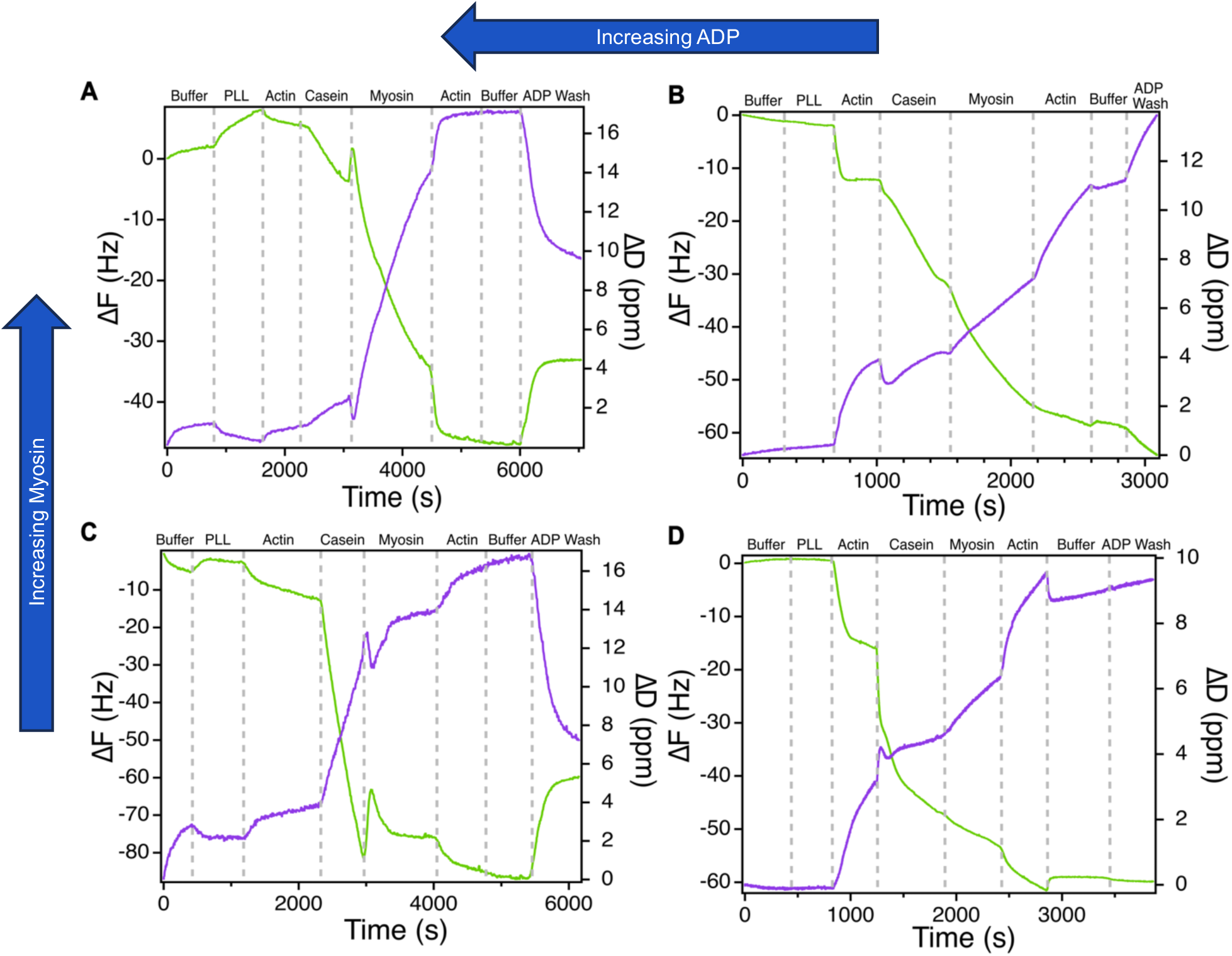
Influences of motor concentration and nucleotide state. Representative QCM-D traces of actomyosin bundle assembly and the influence of modulating myosin II motor concentration and nucleotide binding through final GAB-ADP washes. Frequency changes are shown in green and associated with the left axis, and dissipation changes are shown in purple and associated with the right axis. All buffer washes with 0.2 mM ATP, PLL, casein, and actin are the same across all four panels. (A) 0.6 nM myosin in the bundle, 0.2 mM ADP in final wash. (B) 0.6 nM myosin in the bundle, 0.02 mM ADP in the final wash. (C) 0.3 nM myosin in the bundle, 0.2 mM ADP in the final wash. (D) 0.3 nM myosin in the bundle, 0.02 mM ADP in the final wash. For all traces with 0.6 nM myosin: N=7 with frequency shifts of (all in Hz, average shift ± SEM): buffer 0.7 ± 0.2, PLL 1.7 ± 0.6, first actin 11 ± 1.3, casein 22 ± 3.3, myosin 32 ± 9.0, second actin 10 ± 2.6, ATP wash 2.0 ± 0.6, ADP wash 5.8 ± 2.8; and with dissipation shifts of (all in ppm, average shift ± SEM): buffer 0.2 ± 0.08, PLL 0.2 ± 0.06, first actin 4.1 ± 1.3, casein 1.9 ± 0.5, myosin 8.0 ± 2.7, second actin 4.4 ± 1.4, ATP wash 0.8 ± 0.5, ADP wash 3.2 ± 1.2. N=6 for ATP wash and N=4 for ADP wash. For all traces with 0.3 nM myosin, N=8 with frequency shifts of (all in Hz, average shift ± SEM): buffer 1.4 ± 0.5, PLL 4.8 ± 2.3, first actin 9.4 ± 3.8, casein 31 ± 5.1, myosin 9.8 ± 2.5, second actin 5.8 ± 1.5, ATP wash 4.3 ± 1.8, ADP wash 7.7 ± 3.4, and dissipation shifts of (all in ppm, average shift ± SEM): buffer 1.4 ± 0.5, PLL 1.9 ± 1.0, first actin 3.8 ± 1.6, casein 4.4 ± 1.1, myosin 4.2 ± 1.0, second actin 2.1 ± 0.4, ATP wash 2.0 ± 0.9, ADP wash 4.9 ± 2.3. N=6 for ADP wash. In (A) and (C), a higher concentration ADP buffer wash results in significant frequency increase and dissipation decrease. Both frequency and dissipation shifts here have similar amplitude regardless of myosin concentration. In (B) and (D), lowering the ADP concentration in GAB to 10% of what was used in (A) and (C) yields slight decreases in frequency and increase in dissipation, both of which are commensurate with the respective myosin concentration.

The buffer washes with different nucleotides also yielded unique signatures in frequency and dissipation shifts. In Figure 3, both “buffer” washes are with general actin buffer (GAB) that is supplemented with ATP. Further, all additions of PLL, actin, casein, and myosin are all diluted in GAB. In Figure 3A, frequency and dissipation shift very little with the second ATP buffer addition, which would be expected since we are not adding any substantial mass. In Figures 3B-3D, however, dissipation shifts slightly initially and then levels out. This wash would be removing any remaining unbound actin from the previous step, as well as introducing fresh ATP, and thus an increase in motor activity in the actomyosin bundles. The slight variability seen between the traces arises from the variability in distribution of the elements as they selectively bind their targets on the sensor surface. The increase in dissipation shift for the ATP buffer wash indicates an increase in system softness, suggesting that increasing motor activity alters the viscoelasticity of the local actomyosin system. This correlates with results from Koenderink *et al.* where myosin activity in AF meshes leads to increased viscous dissipation in comparison to a passive actin solution.^33^

A final buffer wash is performed where the same GAB buffer is used except ADP is substituted for ATP. In Figures 3A and 3C, the ADP concentration is the same as the ATP concentration used in the GAB buffer throughout the experiments (0.2 mM). In Figures 3B and 3D, the ADP concentration is one-tenth of that in 3A and 3C. In Figures 3A and 3B, the myosin concertation is double that of 3C and 3D. All other infusion conditions are the same across all four panels. These changes in ADP concentration yield strikingly different trace profiles. In Figures 3A and 3C, the higher concentration ADP wash results in a significant frequency increase and dissipation decrease. The addition of ADP halts myosin activity and is likely seizing up the actomyosin system, leading to an increase in rigidity in comparison to when the motors were active in the presence of ATP. Similar trends have been demonstrated with large-scale AF mesh measurements.^24,33^ The frequency increase is interesting in that we typically interpret this as mass being removed from the sensor surface due to the microfluidic system. However, performing a wash step with GAB containing ATP just before did not affect the frequency significantly. We refer to “mass addition” when the sensor’s vibrations are being dampened. In the case of Figures 3A and 3C, seizing up myosin activity and increasing myosin’s affinity for actin somehow reduces the dampening effect of the actomyosin system on the sensor surface, perhaps through more complex bundling that is not necessarily surface-bound but formed hierarchically away from the surface. In addition, the changes in frequency and dissipation in Figures 3A and 3C are quite similar even though myosin concentration is halved in 3C. This may indicate that the ADP wash affects the actin filaments as well. Previous studies have shown that different nucleotides may alter actin filament stiffness, although there is some debate as to the mechanism.^6,15,25,27,47^

Lowering the ADP concentration results in a different trend from 3A and 3C. In Figure 3B, the 0.02 mM ADP wash yields a slight decrease in frequency and increase in dissipation, and the same trend is in Figure 3D except the magnitudes are smaller (see also Figure S2). Here, the frequency and dissipation changes are commensurate with the respective myosin concentration, indicating that the lower ADP concentration is likely primarily affecting the myosin activity. The lower ADP concentration may also be mixing with some of the previous ATP, making a hybrid nucleotide mixture where some myosins lose activity due to ADP binding and some others remain active. Myosin inactivity and increasing actin affinity due to ADP binding will create transient crosslinks while other myosins remain active, and this may facilitate better coordination and traction for myosin force generation, increase system productivity, and result in increasing dissipation/system compliancy.

There is not a clear precedent for QCM-D data presentation and interpretation for soft matter systems outside of presenting ΔF and ΔD plots, especially in regards to dissipation change due to protein activity as few studies exist, few report dissipation curves, and those that have dissipation curves interpret them qualitatively.^5,31,46,58^ Here, we present the frequency and dissipation shift magnitudes for a particular infusion condition across all experiments in Table 1. The N values in terms of the number of reported QCM-D traces exceed other QCM-D studies that report small (3-5) or even one experimental trace per condition as QCM-D has traditionally been used as a characterization technique.^31^ One of the few *in vitro* cytoskeletal motor investigation examples includes Ozeki *et al.* where they have N=3 for kinesin binding casein and Persson *et al.* who reports an N of 4-5 experiments for binding heavy meromyosin to a sensor surface.^45,46^ However, we note that these experimental examples consisted of only one or two infusion conditions, not the seven or eight infusion conditions per experiment used in this study. Thus, the number of infusion steps, reagent concentration and volume, time of experiments, and similarity between traces justify the Ns presented here.

**Table 1.**
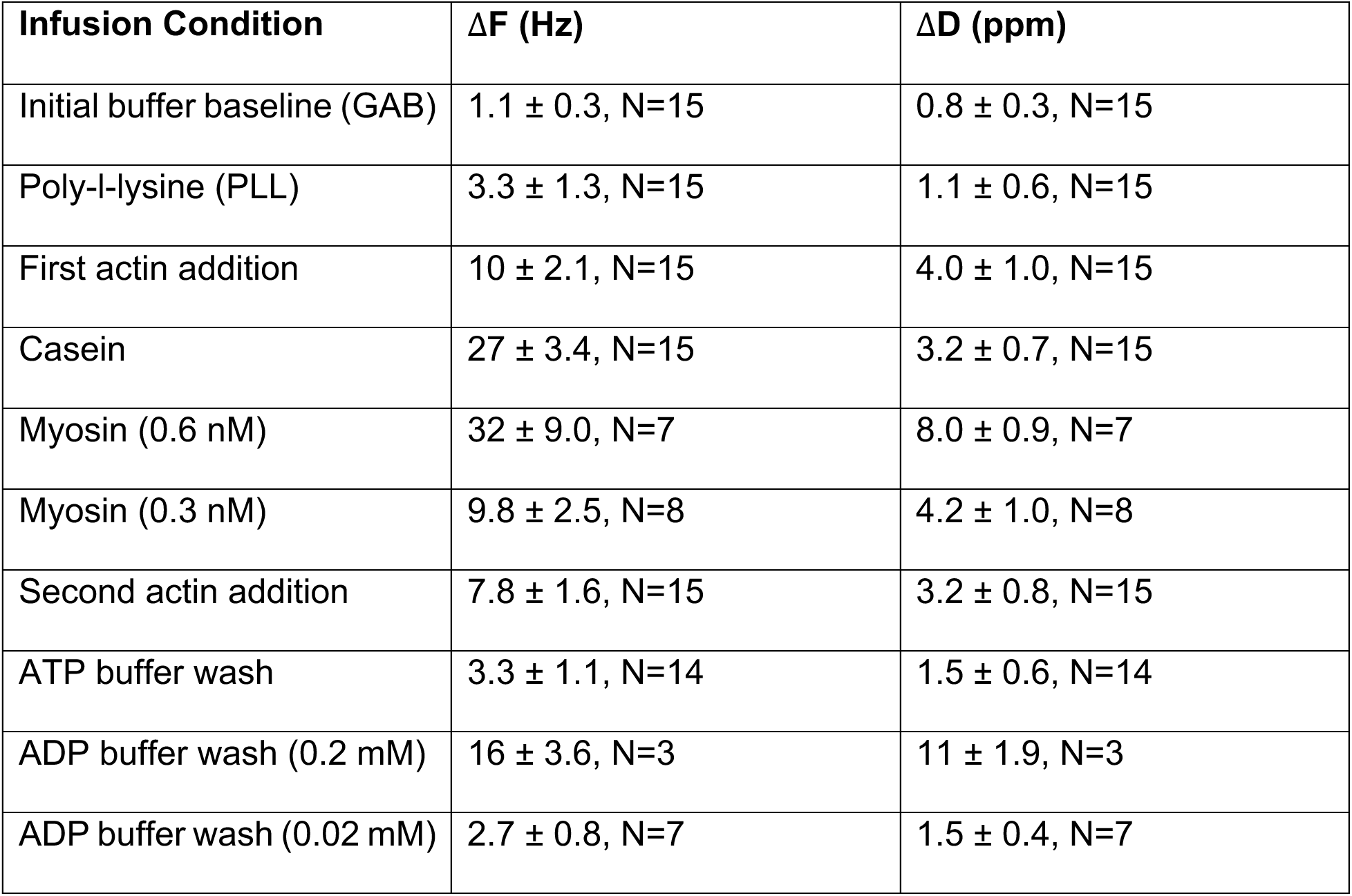
Average magnitude of QCM-D frequency and dissipation shifts in actomyosin bundles. Average ± SEM for each infusion condition collated across experimental runs.

Overall, the average magnitudes of the frequency and dissipation shifts for each infusion condition demonstrate the capabilities of using QCM-D to investigate molecular-level changes to cytoskeletal ensembles. The initial buffer baseline shows very little change in frequency and dissipation, as it should since nothing should be binding the sensor surface. The addition of poly-l-lysine adds mass with negligible change in dissipation, and casein clearly adds mass but little change in dissipation. However, actin and myosin have higher changes in dissipation due to their size and more compliant or “softer” nature. Interestingly, halving the myosin concentration has a concomitant change in ΔF, as expected, but ΔD is slightly closer in magnitude between the two myosin concentration conditions. This may be due to the active nature of myosin, their transient interactions with actin, and their non-linear emergent ensemble mechanics as myosin group size changes that make mechanical changes more distributed throughout the actomyosin system. In addition, the second actin infusion, although at the same concentration as the initial infusion, has smaller magnitudes of ΔF and ΔD. This is likely attributed to the reduced number of binding sites available for the second actin infusion as the actin should only be attaching to myosin that are bound to surface-bound actin filaments. The ATP buffer washes also see minimal changes in frequency and dissipation, but the frequency and dissipation shift direction and magnitude for the ADP washes are highly dependent on ADP concentration. Each infusion condition has a characteristic signature in terms of frequency and dissipation change measurements relative to preceding infusions, but there are also signature features in the trace shape indicating how much time is required for binding and settling onto the existing proteins on the sensor surface. These are all parameters that can be extracted and further explored to better understand the capabilities of the technique and how to interpret the results either alone or in tandem with other techniques.

The actomyosin bundle QCM-D results here are complementary to our previous force generation changes measured by optical tweezers^2^. Myosin II group dynamics, such as stepping and force generation, are significantly different depending on the local environment in which they were working. In addition, it is known that myosin II as a single molecule is non-processive, yet when in groups, they processively work together. But how do the motors know whether to be in a low or high duty ratio mode? We hypothesize that there are two levels of sensory-force feedback: motor-motor communication and more systemically through the AF.^3^ Contributions could be due to myosins existing in a semi-diffusive state with the ability to entropically expand (directional ensemble movement due to diffusion) with some level of force contribution.^3,34^ It may also be due to the number of motors that are attached, as saturation of binding sites would require metachronal wave-like systematic communication in order for the myosins to be able to take forward steps without wasting energy.^41^ In addition, the number of attached motors will change the mechanical properties of the AF (e.g. more motors attached increases AF stiffness) but may also influence pre-stress on the AF.^16,33,37^ Thus, emergent AF compliancy may communicate to distal motors which duty ratio mode is needed for the function at hand.^28,55^ Through the QCM-D results presented, we observe such changes in mechanical compliancy of *in vitro* actomyosin systems due to changes in motor activity and concentration, further backing the hypothesis that actin filaments can serve as a force sensor and mediator of feedback to its bound motors and local environment.^3,8,28,40,55^

The ability of QCM-D to detect molecular level changes in mass and viscoelasticity in cytoskeletal ensemble systems will open doors for a variety of applications and experiments that complement existing techniques, such as total internal reflection fluorescence microscopy, atomic force microscopy, and optical trapping. Further assay customizations could be explored, such as using different sensor cleaning protocols, coatings, and surface modifications. For instance, one could use silicon dioxide sensors to more closely mimic coverslips used for microscopy experiments or buy protein-coated sensors, such as biotin. Experiments could also be expanded to other cytoskeletal systems and motor proteins as the addition of environmental influences, such as crowding agents, ionic strength, and crosslinkers have been suggested to influence motor behavior.^26,28,36,43,55^ For example, the effect of flexible crosslinkers like filamin A alone or myosin II motors alone in AF meshes have marginal influences on system stiffening, yet when combined, have significant emergent effects on stiffness and activity.^33^ Increasing the complexity of these *in vitro* ensembles will approach more physiologically relevant conditions to then compare how their systemic viscoelasticity changes correlate with motility or force generation changes in TIRF or optical trapping experiments, respectively. However, there are pitfalls that remain in probing cytoskeletal assemblies using QCM-D. For instance, the QSense Analyzer has included software that will provide estimates of Young’s modulus, coating thickness, and mass added to the sensor. One of the major assumptions of the Sauerbrey equation used for detecting mass and then ultimately obtaining estimates for modulus is that the sensor has a complete, homogenous, and rigid coating of even thickness. That is certainly not the case with this particular application as we are using multiple proteins of different sizes and compliances, but what likely has more influence is the non-uniformity of the “coating” thickness, as schematically shown in Figure 1B. Perhaps this would be less of an issue when exploring large-scale AF meshes, but when attempting to characterize small motor ensemble properties, the sensor is not evenly coated. Thus, correction factors or different models would need to be developed and adapted with the software in order to use them for quantitative analysis. However, with strict controls and notes of when design parameters are changed, the real-time measurement of frequency and dissipation changes within cytoskeletal systems can be compared amongst each other to determine how potential mechanical influencers, such as different protein types, environmental buffer conditions, or nucleotide state, alter the cytoskeletal ensemble’s mechanics. Other potential downsides to these experiments are the high concentrations of protein required and the time required per experiment with our longest runs being almost two hours of data, but that does not include the fresh protein solution preparations and sensor and QCM module cleaning required before and after use. One should also consider how QCM-D data would supplement their specific mechanistic inquiry. Until improved models are developed for quantitatively interpreting QCM-D frequency and dissipation change measurements for cytoskeletal ensembles, it would be beneficial to couple the qualitative information regarding changes in assay conditions and system viscoelasticity with quantitative force measurement techniques, like AFM or optical tweezers, or in conjunction with fluorescence microscopy.^31^ For example, Yang *et al.* characterized cell-level mechanical behavior of epithelial cells in response to epidermal growth factor (EGF) using a combination of fluorescence imaging to capture the response of actin filaments, AFM to probe the cell’s mechanical response, and QCM-D to correlate energy with the measured AFM response to EGF stimulation.^61^ Their approach demonstrates the advantage of using complementary techniques to formulate a more complete biophysical profile of their system of interest and should be applied more readily to *in vitro* studies.

In conclusion, we demonstrate that QCM-D is a useful tool that complements existing experimental approaches in systematically characterizing the interconnection of force, mechanics, and machinery within cytoskeletal ensembles and can be used to decipher which mechanical cues lead to certain ensemble behaviors.^35^ QCM-D will especially be helpful in constructing more thorough biophysical profiles of cytoskeletal ensembles through identifying the molecular parameters that dictate emergent mechanical behavior and revealing clues towards the design principles of regulatory feedback behavior at the molecular level that then propagates up in scale to influence long-distance cellular tasks.

## Acknowledgements

This work is supported by the American Heart Association #848586, NIH R35 GM147050, NSF REU Site: Ole Miss Nanoengineering Summer REU Program NSF ENG/EEC #2148764, and the Mississippi Space Grant Consortium under grant number NNX15AH78H. QCM-D instrumentation, data, and analysis are supported by NSF MRI grant number 2018004. The shared QCM-D facility at the University of Mississippi is open to outside collaboration. If interested, contact.

## Author Contributions

EMK, VNA, and DNR conceptualized the work. EMK, VNA, WBW, LHH, LT, and OYA developed and optimized the protocols. EMK, VNA, WBW, and LHH gathered and analyzed the data. EMK, VNA, and DNR wrote and edited the manuscript.

## Supplemental Information

Detailed methods and additional QCM-D traces can be found in the supplemental information. sample

