## Supplemental Methods and Figures for "Deciphering Mechanochemical Influences of Emergent Actomyosin Crosstalk using QCM-D"

**QCM-D Instrument**

The QSense Analyzer from Biolin Scientific was used for all experiments, and Biolin's Dfind software was used for data acquisition. The standard sensor module was used at a temperature of 25°C for all experiments.

**QCM Sensor Preparation**

QCM sensors must be cleaned before and after data collection. Gold-coated QSense sensors (QSX 301, Biolin Scientific) were used for all experiments. To clean the sensors before use, the module was set to heat to 25-30°C, and the peristaltic pump was set to 0.1-0.3 mL/min. For each module being used, 5 mL of 2% Hellmanex was added to the module through the pump and microfluidic system. During this step, the max flow button was occasionally pressed to ensure cleaning of the surface. This step was repeated with deionized water. Once the water completely ran through the module, the inlet tubing was removed from the water and allowed air to flow through the tubing until no liquid could be seen exiting the tubing. The sensor was then carefully removed from the module using tweezers, taking care to not touch the gold surface, and dried with nitrogen gas. The module was then removed from the QCM-D to dry the interior with nitrogen gas. When using sensors for the first time, they were cleaned by placing them in a UV-ozone chamber for 10 minutes. Subsequently, the sensors were then submerged in 2 wt% SDS in deionized water for 30 minutes at room temperature. Using tweezers to hold the sensor, they were then rinsed with deionized water and dried with nitrogen gas. The sensors were placed back into the UV-ozone chamber for 10 minutes. The sensors were then immediately placed into the QCM module for experiments.

For sensors that are reused after experiments, it is recommended to use piranha solution for cleaning to ensure that all adhered protein remnants are removed. To use this method, create a 5:1:1 solution of reverse osmosis (RO) water, 25% ammonia, and 30% hydrogen peroxide. In a fume hood, heat 20 mL of RO water to 80°C in a 100 mL beaker. After the temperature is reached, add 4 mL of 25% ammonia and 4 mL of 30% hydrogen peroxide. As the temperature adjusts back to 80°C, carefully place sensors into a dipping rack. Place the stand into the beaker and allow the sensors to remain the solution for 5-10 minutes, or until bubbles appear on the sensors. Remove the sensor dipping rack and rinse each sensor individually with RO water. Immediately dry with nitrogen or air. Store the sensor in an enclosed container to avoid contamination before measurements.

**Actin Polymerization**

Actin polymerization was performed as described previously.[1–4] Non-labeled rabbit skeletal muscle actin (Cytoskeleton) was reconstituted by adding 100 μL of reverse osmosis (RO) water to 1 mg of lyophilized actin. The contents were mixed by gently pipetting up and down, aliquoted, and stored at -80°C with final actin concentration of 10 mg/mL. To polymerize non-labeled actin into filaments, 5 μL of 10 mg/mL actin were mixed with 100 μL General Actin Buffer (GAB: 5 mM Tris-HCl, 0.2 mM $CaCl_2$, 0.5 mM DTT, 0.2 mM ATP). The mixture was kept on ice and allowed to incubate for one hour. Actin was then polymerized into filaments by adding 5.5 μL of Actin Polymerizing Buffer (APB: 50 mM Tris-HCl, 500 mM KCl, 2 mM $MgCl_2$, 2 mM $CaCl_2$ 2 mM DTT, 5 mM ATP) to the actin mixture, mixed well by gently pipetting up and down, and allowed to incubate for 20 minutes on ice. Actin filaments were stabilized and fluorescently labeled by adding 5 μL of rhodamine-labeled phalloidin (Cytoskeleton). The vial was wrapped in aluminum foil to block light and allowed to incubate on ice for one hour. The mixture was stored at 4°C to be used for preparing actin myosin bundles for up to one week.

**Actomyosin Bundle QCM-D Experiments**

Solutions for actomyosin QCM-D experiments include: General Actin Buffer (GAB): 5 mM Tris-HCl, 0.2 mM $CaCl_2$, 0.5 mM DTT, 0.2 mM ATP; Poly-l-lysine (PLL): 0.1 w/v% PLL in water (Sigma Aldrich) further diluted by adding 30 μL into 3 mL GAB; Actin filaments (AF): 2 μL of concentrated stock actin solution per mL of GAB; Casein: Blotting-grade blocker (Bio-rad), 1 mg per 15 mL GAB; Skeletal myosin II: Full-length skeletal myosin II (Cytoskeleton) at 0.6 nM in GAB (or 0.3 nM for lower concentration experiments); and GAB-ADP: 5 mM Tris-HCl, 0.2 mM $CaCl_2$, 0.5 mM DTT, 0.2 mM ADP (or 0.02 mM ADP for lower concentration experiments).

A cleaned gold sensor is placed into the QCM module to prepare for component addition. The peristaltic pump is set to 0.1 mL/min, and the module temperature is set to 25°C. Each solution is kept on ice in conical 15 mL tubes until ready for use when the QCM tubing is submerged into the respective solution for addition to the sensor. First, GAB is added as the baseline solution for 10-15 minutes until the frequency signal plateaus. Then, the sample tubing will be switched to PLL, actin, casein, myosin, actin, and GAB in subsequent steps after the previous addition plateaus or before the sample runs out. There is generally a span of a few minutes between switching the inlet tubing to a new sample and observing frequency changes on the computer.

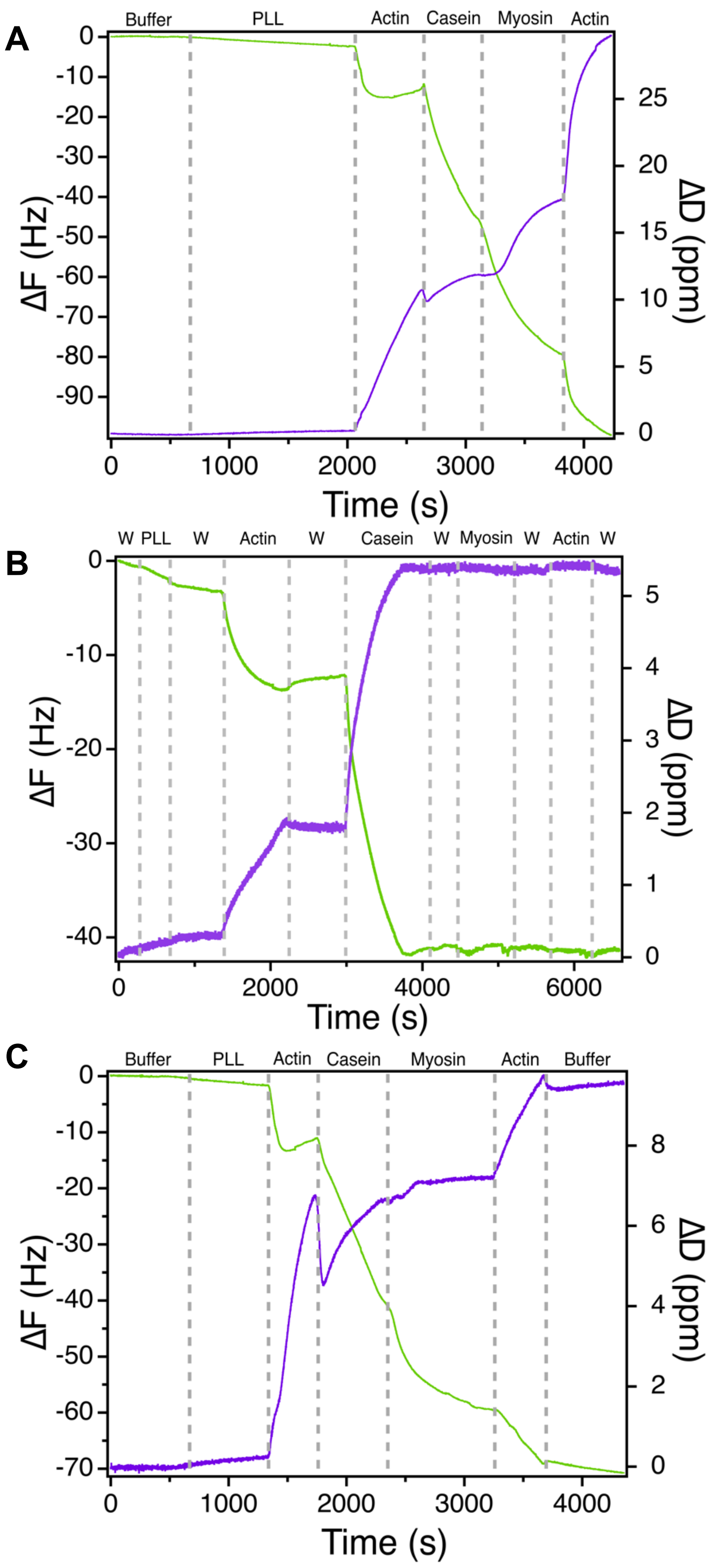


**Supplemental Figure S1 | Determining Conditions for Actomyosin Activity and Sensitivity using QCM-D.** Representative traces of assay conditions tested to understand how sensitive changes in actomyosin bundle conditions will be represented by QCM-D. (A) Actomyosin bundle formation with no wash steps. The poly-l-lysine addition is longer than in other traces in an attempt to see how long it would take to for the signal to stabilize. (B) Wash steps of ATP-based GAB buffer were incorporated after every addition to evaluate if the washes would aid or inhibit assay conditions or sensitivity. The washes not only added significant time to an already lengthy experiment, but they seemed to inhibit subsequent signal sensitivity, especially after the casein addition. It is possible that since the ATP wash buffer was not supplemented with casein that it was actually washing away the existing layer of blocking so myosin motors were transiently sticking to not just the actin filaments, but all over the sensor, thus diluting myosin binding and signal sensitivity. (C) For comparison, only the 7$^{th}$ harmonic from Figure 2 in the manuscript is shown with a final ATP buffer wash. After the entire bundle is formed, the buffer wash has little to no effect on the signal, indicating that the bundles have formed completely and are not washing away.

**Supplemental Figure S2 | Effect of myosin II nucleotide state and concentration on QCM-D signal and bundle viscoelasticity.** QCM-D traces that incorporate a final ATP buffer wash followed by a 0.02 mM ADP buffer wash. Change in myosin nucleotide state from ATP to a lower ADP concentration yields an increase in ΔD, which indicates the surface "coating" is less rigid. This is in contrast to using 1:1 substitution of ADP for ATP, as shown in Figure 3. There is a slight increase in ΔF as well, but not on a commensurate level as the change in dissipation. (A) and (B) assay conditions were the same where the myosin concentration is twice that of in (C). The initial actin ΔF signal is higher in (B) than in (A) (18 vs. 12 Hz), which seems to have allowed more myosin to bind actin in (B). However, there was also less casein binding in (B), which could also contribute to additional myosin binding. This is an example of the variability that can be present in QCM-D experiments. While the absolute values of ΔF and ΔD may slightly differ between traces under the same assay conditions, major assay condition changes, such as change in concentration and nucleotide state, are still evident. (C) A reduction in myosin binding due to using half the concentration is evident in the overall decrease in ΔF from (A) or (B). Commensurate myosin presence and activity in each trace is also captured in the ADP wash ΔD signal. More myosin present in the actomyosin bundles yields a greater effect in the change in nucleotide state from the weakly bound ATP state to the strongly bound ADP state.

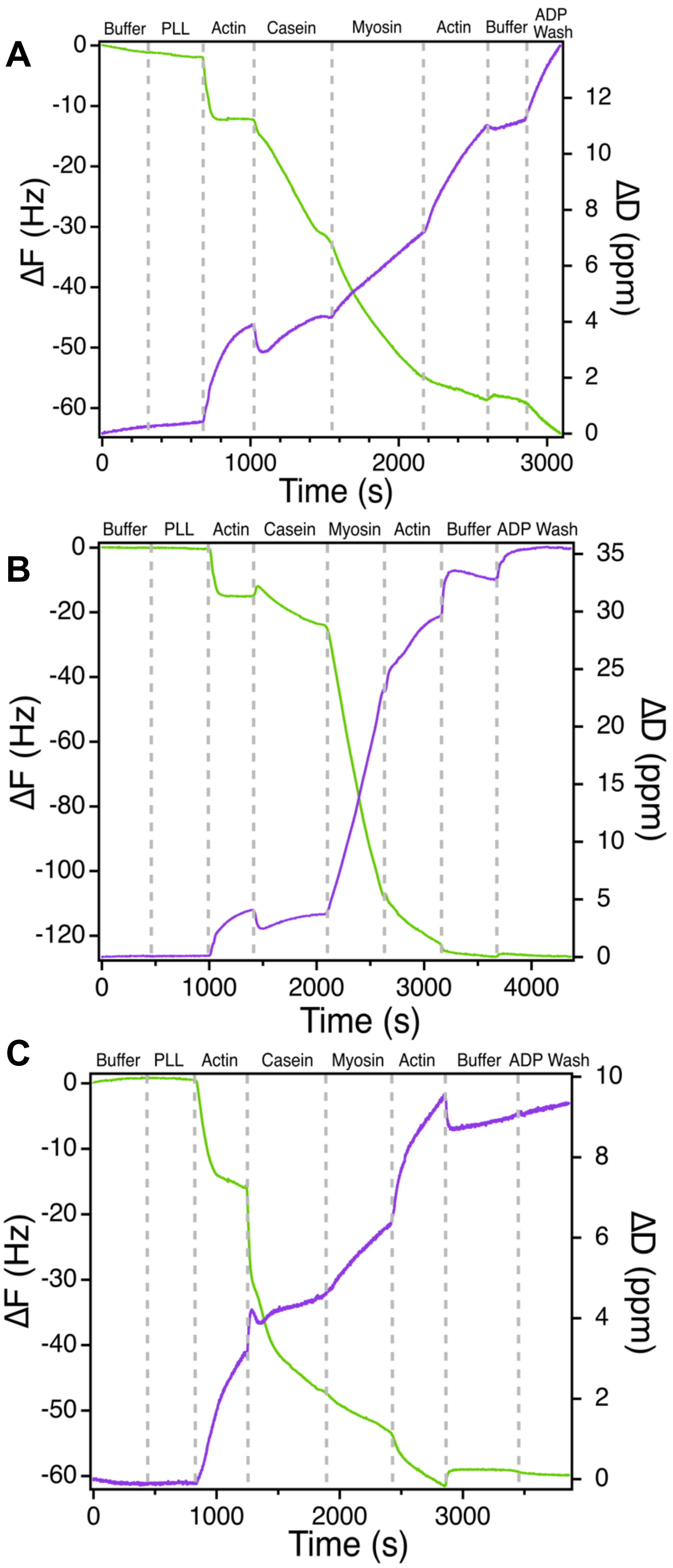
